# Context-dependent activation of SIRT3 is necessary for anchorage-independent survival and metastasis of ovarian cancer cells

**DOI:** 10.1101/670778

**Authors:** Yeon Soo Kim, Piyushi Gupta-Vallur, Victoria M. Jones, Beth L. Worley, Sara Shimko, Dong-Hui Shin, LaTaijah C. Crawford, Chi-Wei Chen, Katherine M. Aird, Thomas Abraham, Trevor G. Shepherd, Joshua I. Warrick, Nam Y. Lee, Rebecca Phaeton, Karthikeyan Mythreye, Nadine Hempel

## Abstract

Cells must alter their antioxidant capacity for maximal metastatic potential. However, the antioxidant adaptations required for transcoelomic metastasis, which is the passive dissemination of cancer cells in the peritoneal cavity as seen in ovarian cancer, have largely remained unexplored. Contradicting the need for oxidant scavenging by tumor cells is the observation that expression of the nutrient stress sensor and regulator of mitochondrial antioxidant defenses, SIRT3, is suppressed in many primary tumors. We discovered that this mitochondrial deacetylase is however, upregulated in a context-dependent manner in cancer cells. SIRT3 activity and expression transiently increased following ovarian cancer cell detachment and in tumor cells derived from malignant ascites of high-grade serous adenocarcinoma patients. Mechanistically, SIRT3 prevents mitochondrial superoxide surges in detached cells by regulating the manganese superoxide dismutase SOD2. This mitochondrial stress response is under dual regulation by SIRT3. SIRT3 rapidly increases SOD2 activity as an early adaptation to cellular detachment, which is followed by SIRT3-dependent transcriptional increases in SOD2 during sustained anchorage-independence. In addition, SIRT3 inhibits glycolytic capacity in anchorage-independent cells thereby contributing to metabolic changes in response to detachment. While manipulation of SIRT3 expression has few deleterious effects on cancer cells in attached conditions, SIRT3 up-regulation and SIRT3-mediated oxidant scavenging following matrix detachment are required for anoikis resistance *in vitro*, and both SIRT3 and SOD2 are necessary for colonization of the peritoneal cavity *in vivo*. Our results highlight the novel context-specific, pro-metastatic role of SIRT3 in ovarian cancer.

## Introduction

Epithelial ovarian cancer (EOC) remains the most deadly gynecological malignancy, with a five year survival rate of less than 29% for patients diagnosed with advanced stage metastatic disease [1]. During transcoelomic spread the peritoneal fluid facilitates the dissemination of detached ovarian cancer cells to peritoneal cavity organs, including the omentum [2, 3]. This is evident by malignant ascites accumulation in most advanced stage patients [3]. Thus evasion of anchorage-independent cell death (anoikis) upon detachment from the primary tumor is a likely critical step for ovarian cancer metastasis [4]. A feature of anoikis is the surge in oxidative stress elicited by cell detachment. Adaptations to oxidative stress are therefore necessary for successful hematological metastatic spread, as demonstrated in melanoma and breast cancer [5–7]. Moreover, administration of compounds that facilitate oxidant scavenging, such as N-acetyl-cysteine, can promote metastasis *in vivo* [8]. However, it remains largely unexplored if adaptations to oxidative stress are required by ovarian cancer cells for successful transcoelomic metastasis.

Contradicting the need of tumor cells for oxidant scavenging is the observation that expression of the nutrient stress sensor and regulator of mitochondrial antioxidant defenses, the Sirtuin deacetylase SIRT3 [9–12], is suppressed in many primary tumors [13–17]. Moreover, several studies have demonstrated that SIRT3 knock-down promotes proliferation and tumorigenesis in tumor models of breast [12, 18], mantle cell lymphoma [19] and liver cancer [16], promoting investigators to initially characterize SIRT3 as a tumor suppressor. However, it is becoming increasingly clear that the role of SIRT3 in tumor biology is complex [17, 20, 21]. Pro-tumorigenic properties of SIRT3 have conversely been reported in oral squamous cell carcinoma [22], and colorectal cancer [23], with increased SIRT3 expression being associated with poor outcome in colon and non-small cell lung cancer patients [17]. In addition, SIRT3 promotes glioblastoma multiforme (GBM) stem cell viability [24], and is an important component of the mitochondrial unfolded protein response (mtUPR) necessary for breast cancer metastasis [25]. The latter function of SIRT3 is being attributed to its role as a regulator of the antioxidant response required for tumor cell survival and metastasis.

Although, previous reports have demonstrated that SIRT3 exerts anti-proliferative and anti-migratory effects on ovarian cancer cells [26, 27], the role of SIRT3 during ovarian cancer transcoelomic spread has not been investigated. Moreover, when and where SIRT3 is expressed during tumor progression remains unknown. We discovered that SIRT3 is upregulated in a context-dependent manner in ovarian cancer cells, and indeed has a specific pro-metastatic role, by supporting anchorage-independent survival. While SIRT3 expression is low in primary ovarian tumors and knock-down of its expression has no deleterious consequences in attached proliferating conditions, we demonstrate that SIRT3 activity and transcription are specifically induced in response to anchorage-independence, and that this transient increase results in the activation of the mitochondrial antioxidant SOD2, which is necessary for anchorage-independent survival and peritoneal colonization *in vivo*. These findings provide important evidence of the function of SIRT3 in ovarian cancer, and clarify several contradictory findings associated with the role and expression of SIRT3 in cancer progression.

## Results

### SIRT3 expression increases in a context-dependent manner during ovarian cancer metastasis and is induced in response to matrix detachment

Expression analysis of primary ovarian tumors and matching cells derived from malignant ascites, and omental and peritoneal metastatic lesions from a publicly available data set (GEO:GSE85296) revealed that SIRT3 levels are heterogeneous in tissues representing different stages of transcolomic metastatic spread (**Fig. 1A**). While SIRT3 levels were lowest in primary ovarian tumors, highest SIRT3 expression was found in cells derived from malignant ascites. In two of four patient samples, the increase in SIRT3 expression was maintained in omental and peritoneal metastatic lesions, while in the other cases SIRT3 expression reverted to levels observed in primary tumors of the ovary. These data suggest that detachment induces transient increases in SIRT3 expression in malignant ascites.

**Figure 1:**
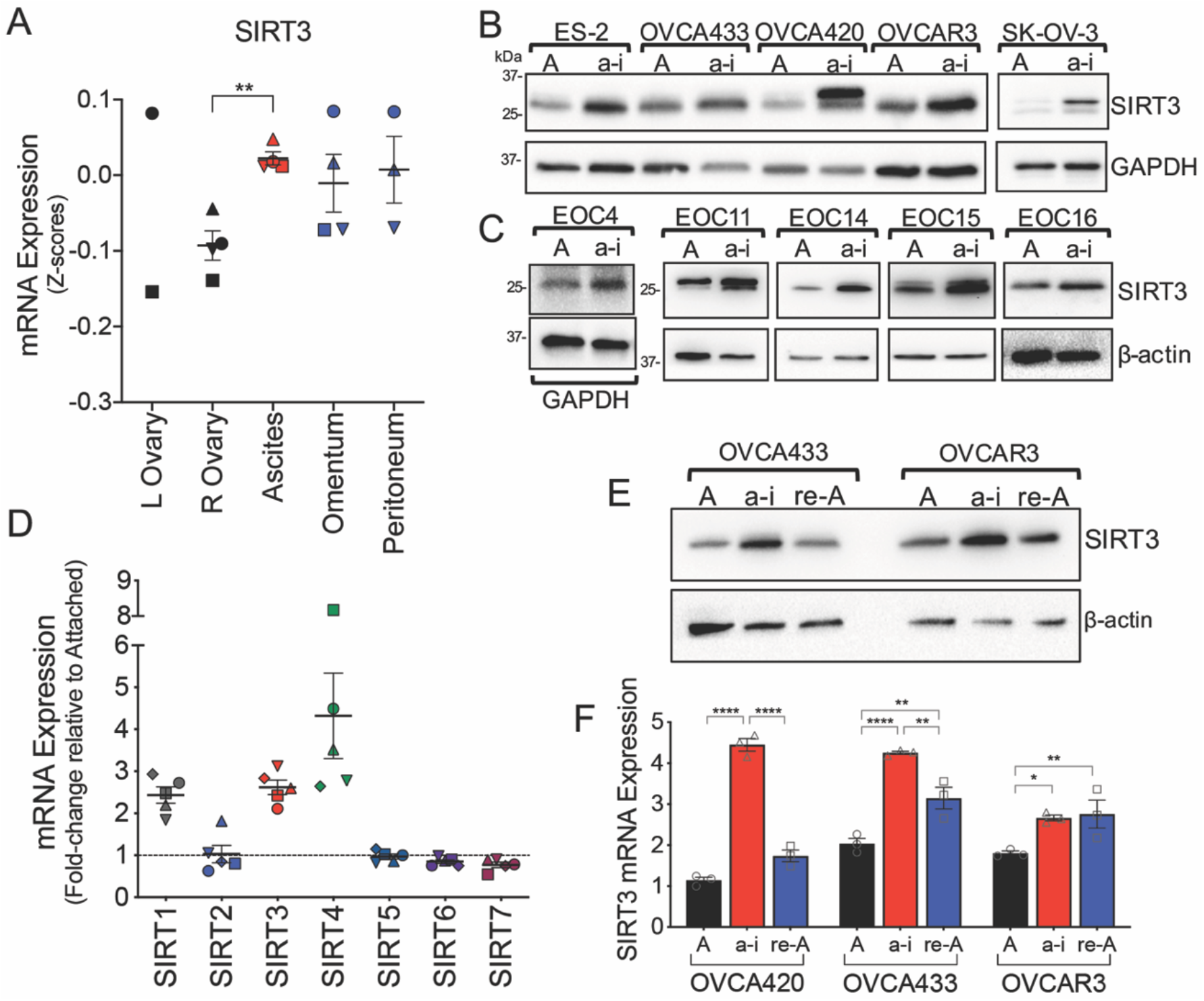
Ovarian cancer SIRT3 expression is context-dependent, and increases in response to anchorage-independence. **A.** SIRT3 mRNA expression in tumor samples from primary tumors of the ovary, matching ascites, and omental and peritoneal metastatic lesions (Geo:GSE85296, n=4, mean expression at each site per patient shown, repeated measures ANOVA P = 0.047, Dunnett’s multiple comparison test **P = 0.005). **B.** Ovarian cancer cell lines were cultured in anchorage-independent (a-i) conditions using ultra low attachment (ULA) plates for 72 h, and SIRT3 protein expression compared to cell cultures in attached (A) conditions using western blotting. **C.** Epithelial ovarian cancer cells (EOCs) were derived from ascites of Stage III and IV high grade serous adenocarcinoma patients and cultured in attached or a-i conditions for 72 h. SIRT3 expression was assessed as in B. **D.** mRNA levels of sirtuin family members were assessed using U133 microarray after EOCs derived from patient ascites (n=5) were cultured in a-i conditions for 72 h as described in [45]. mRNA levels are expressed relative to cells grown in attached conditions. **E.** SIRT3 protein expression was assessed by western blotting of lysates from attached cell cultures (A), cells maintained for 24 h in ULA plates (a-i) and 24 hours following re-attachment (re-A). **F.** SIRT3 mRNA expression was assessed by semi-quantitative real time RT-PCR following cell culturing as in F, and expressed relative to OVCA420 in attached conditions (n=3; two-way ANOVA, Tukey’s multiple comparison test *P<0.05, **P<0.001, ****P<0.0001).

We next tested if a loss of matrix attachment induces SIRT3 expression. Consistently, we found that SIRT3 expression increased when ovarian cancer cell lines (**Fig. 1B**), and ascites-derived primary epithelial ovarian cancer cells (EOCs, **Fig. 1C**) from Stage III and IV high grade serous adenocarcinoma patients were cultured under anchorage-independence in ultra-low attachment (ULA) dishes. RNA expression analysis of EOCs from a separate patient cohort revealed that SIRT3, SIRT1 and SIRT4 were the only members of the sirtuin gene family responsive to mRNA increases when maintained in anchorage-independent conditions (**Fig. 1D**). Interestingly, the transient increase in SIRT3 expression observed in patient ascites (**Fig. 1A**) could be recapitulated in 2/3 cell lines tested, where SIRT3 expression reverted to basal levels following cell re-attachment (**Fig. 1E&F**). These data highlight that SIRT3 expression in ovarian cancer is transient, and induced in a context-dependent manner in response to matrix detachment.

### SIRT3 regulates mitochondrial reactive oxygen species scavenging by activating SOD2 in anchorage-independence

Matrix detachment elevates cytosolic [6] and mitochondrial reactive oxygen species [28], which are thought to contribute to anoikis of non-transformed epithelial cells [6]. To determine if SIRT3 protects cells from mitochondrial redox stress in anchorage-independence, we monitored MitoSox fluorescence in response to SIRT3 knock-down, using several independent si/shRNAs (**Fig. 2A & Supp. Fig. 1A**). Inhibition of SIRT3 expression increased MitoSox fluorescence in anchorage-independent conditions (**Fig. 2B**), indicating that SIRT3 is necessary to maintain low mitochondrial superoxide (O_2^·^_^-^) levels following detachment. Interestingly, SIRT3 knock-down had no effect on MitoSox fluorescence in attached conditions (**Supp. Fig. 1B**), suggesting that SIRT3 has a specific role in protecting against mitochondria oxidant stress during anchorage-independence.

**Figure 2:**
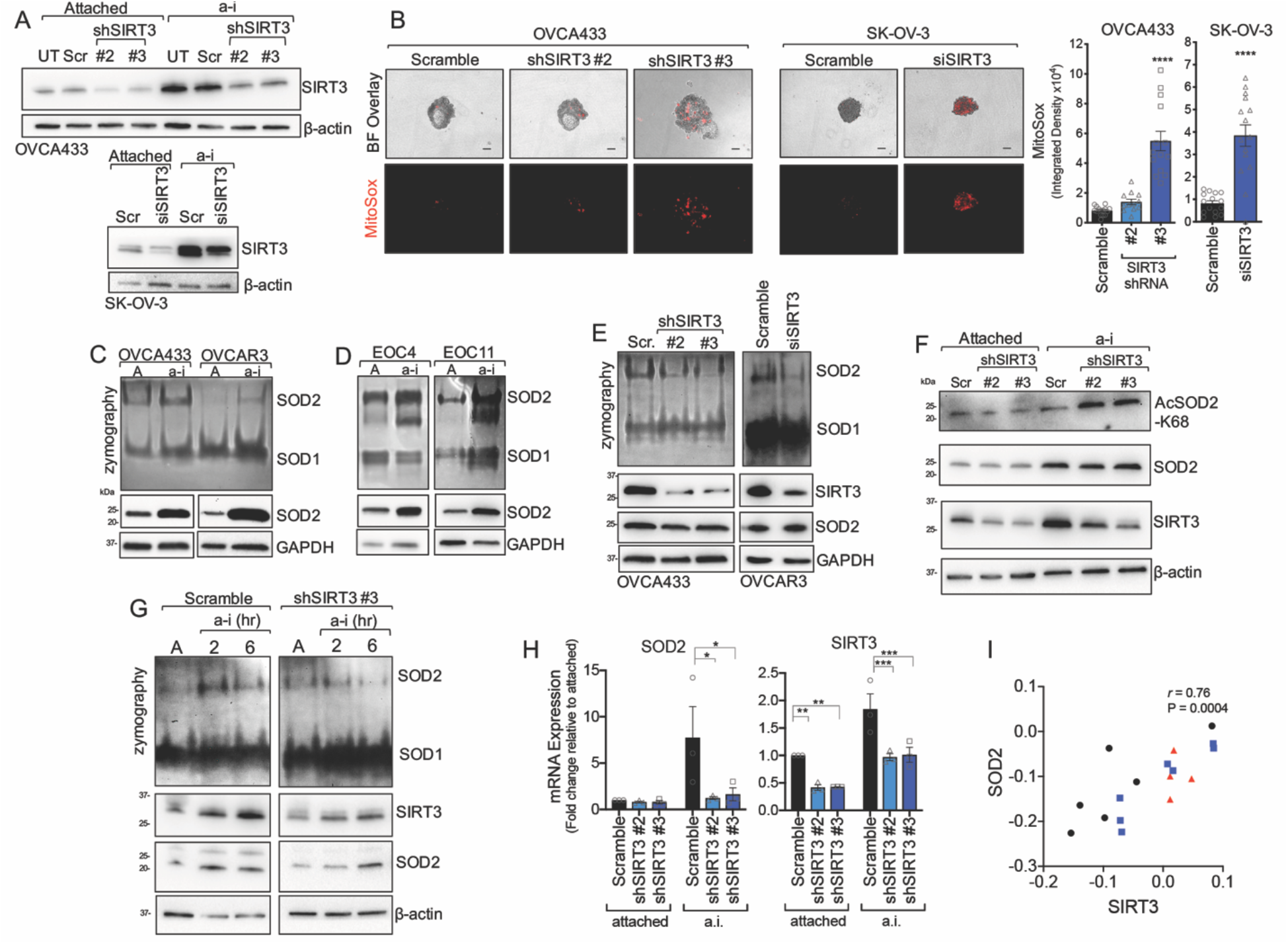
SIRT3 maintains superoxide (O_2^·^_^-^) scavenging in anchorage-independent conditions by activating SOD2. **A.** SIRT3 expression was inhibited using stable shRNA transfection of OVCA433 cells or transient delivery of siRNA in SK-OV-3 (UT, un-transfected; Scr, scramble control). **B.** SIRT3 knock-down increases oxidation and fluorescence of the mitochondrial O_2^·^_^-^ probe MitoSox in ULA cultured OVCA433 (A) and SK-OV-3 (B) ovarian cancer cells. Quantification of MitoSox signal (n=12-14 ± SEM; OVCA433: one-way ANOVA P<0.0001, Tukey’s multiple comparisons test ****P<0.0001, **P=0.001; SK-OV-3: unpaired t-test ****P<0.0001, scale bar = 100μm). **C&D**. Ovarian cancer cell lines (C) and patient ascites-derived epithelial ovarian cancer cells (EOC, D) were cultured in anchorage-independent (a-i) conditions for 72 h using ultra low attachment (ULA) plates. SOD2 activity was assessed by in gel zymography and compared to cell cultures in attached (A) conditions. **E.** shRNA and siRNA-mediated SIRT3 knock-down inhibits SOD2 activity in ULA cultured (72 h) ovarian cancer cells, assessed by SOD zymography. **F.** SIRT3 knock-down increases SOD2 acetylation at lysine 68 in OVCA433 cells cultured in a-i conditions. **G.** SOD2 activity is induced within 2 h of matrix detachment in a SIRT3-dependent manner. SOD2 activity was assessed by zymography in attached (A) OVCA433 cells and cells cultured for 2 or 6 h in anchorage-independence (a-i). **H.** SIRT3 knock-down inhibits SOD2 transcription in a.i.. mRNA expression was assessed by semi-quantitative real time RT-PCR following cell culturing in ULA plates for 24 h. Data expressed relative to expression in scramble transfected cells in attached conditions (n=3; two-way ANOVA, Dunnett’s multiple comparison test *P<0.05, **P<0.01, ***P<0.001). **I.** Positive correlation between SIRT3 and SOD2 mRNA expression in tumor tissues derived from primary ovarian tumors (●), ascites (Δ), and peritoneal or omental lesions (□; Geo:GSE85296, Pearson correlation).

A major antioxidant target of SIRT3 is manganese superoxide dismutase 2 (SOD2), which is one of three superoxide dismutases in the cell, and the primary enzyme responsible for the dismutation of O_2^·^_^-^ to hydrogen peroxide (H_2_O_2_) in the mitochondrial matrix. SIRT3 regulates SOD2 at both the transcriptional level, *via* deacetylaton and activation of the *SOD2* transcription factor FOXO3a [25, 29], and by directly deacetylating and activating SOD2 dismutase activity [9–12]. Concomitant to SIRT3 increases, SOD2 activity and expression were strongly induced in response to detachment of ovarian cancer cell lines (**Fig. 2C**) and patient ascites-derived cells (**Fig. 2D**), indicating that the SIRT3/SOD2 axis is an important adaptation for anchorage-independence.

SIRT3 was directly responsible for enhanced SOD2 activity in detached cells, as evident by SIRT3 sh/siRNA mediated knock-down (**Fig. 2E**). This was accompanied by an increase in SOD2 acetylation at lysine 68, specifically in anchorage-independent conditions (**Fig. 2F**). Importantly, we observed that the SIRT3-dependent increase in SOD2 activity is an early response following matrix detachment. SOD2 activity rapidly increased within 2 hours of matrix detachment, prior to detectable changes in *SOD2* transcription, and this early increase in SOD2 dismutase activity was abrogated by SIRT3 knock-down (**Fig. 2G, Supp. Fig. 1C**).

In addition to this early regulation of SOD2, we observed increased SOD2 expression after 6 hours following matrix detachment, and this was also significantly abrogated by SIRT3 knock-down (**Fig. 2H**). Notably, transcript levels of *SIRT3* in patient specimens from GEO data set GSE85296 strongly correlated with *SOD2* levels, where high expression was similarly observed in ascites-derived cells (**Fig. 2I**). These data demonstrate that the mitochondrial O_2^·^_^-^ scavenger SOD2 is dually regulated by SIRT3 following detachment, and that the SIRT3/SOD2 axis is an early adaptation to anchorage-independence.

### SIRT3 knock-down increases glycolysis in anchorage-independent cells

Given that SIRT3 has previously been associated with suppression of glycolysis [18, 30], we set out to determine if increased SIRT3 expression alters glycolytic flux following cellular detachment. First, we determined if glucose consumption and lactate production are altered in in detached cells, and found that overall glucose consumption was increased, while relative lactate production to glucose consumption was decreased, compared to attached cells (**Fig. 3A**). This suggests that cells in anchorage-independence re-route glucose consumption away from lactic acid production. SIRT3 knock-down significantly increased the ratio of lactate production to glucose consumption in detached cells (**Fig. 3B & Supp. Fig. 2**), and assessment of the optical redox ratio of the metabolic coenzymes FAD and NAD(P)H using multiphoton imaging [31] demonstrated a significant decrease in the FAD / [FAD + NAD(P)H] ratio with SIRT3 knock-down in anchorage-independent cells (**Fig. 3C**). To test if the above results indicate that increased SIRT3 expression switches glucose utilization away from lactate production towards Oxidative Phosphorylation, we examined changes in extracellular acidification rate (ECAR) and oxygen consumption rate (OCR) using extracellular flux analysis. As expected, addition of glucose rapidly increased ECAR in both attached and detached cells (**Fig. 3D**). SIRT3 knock-down significantly increased basal glycolytic rate (basal ECAR) in detached cells following glucose addition, while it had no significant effect on basal ECAR in attached conditions (**Fig. 3E**). While cells in anchorage-independence displayed similar basal glycolytic rate, their maximal glycolytic capacity, stimulated by suppression of respiration with the mitochondrial ATP synthase inhibitor Oligomycin A, was significantly decreased compared to attached cells. Moreover, this was reversed when SIRT3 expression was suppressed (**Fig. 3F**). These data demonstrate that SIRT3 suppresses glycolytic capacity of tumor cells in anchorage-independent conditions.

**Figure 3:**
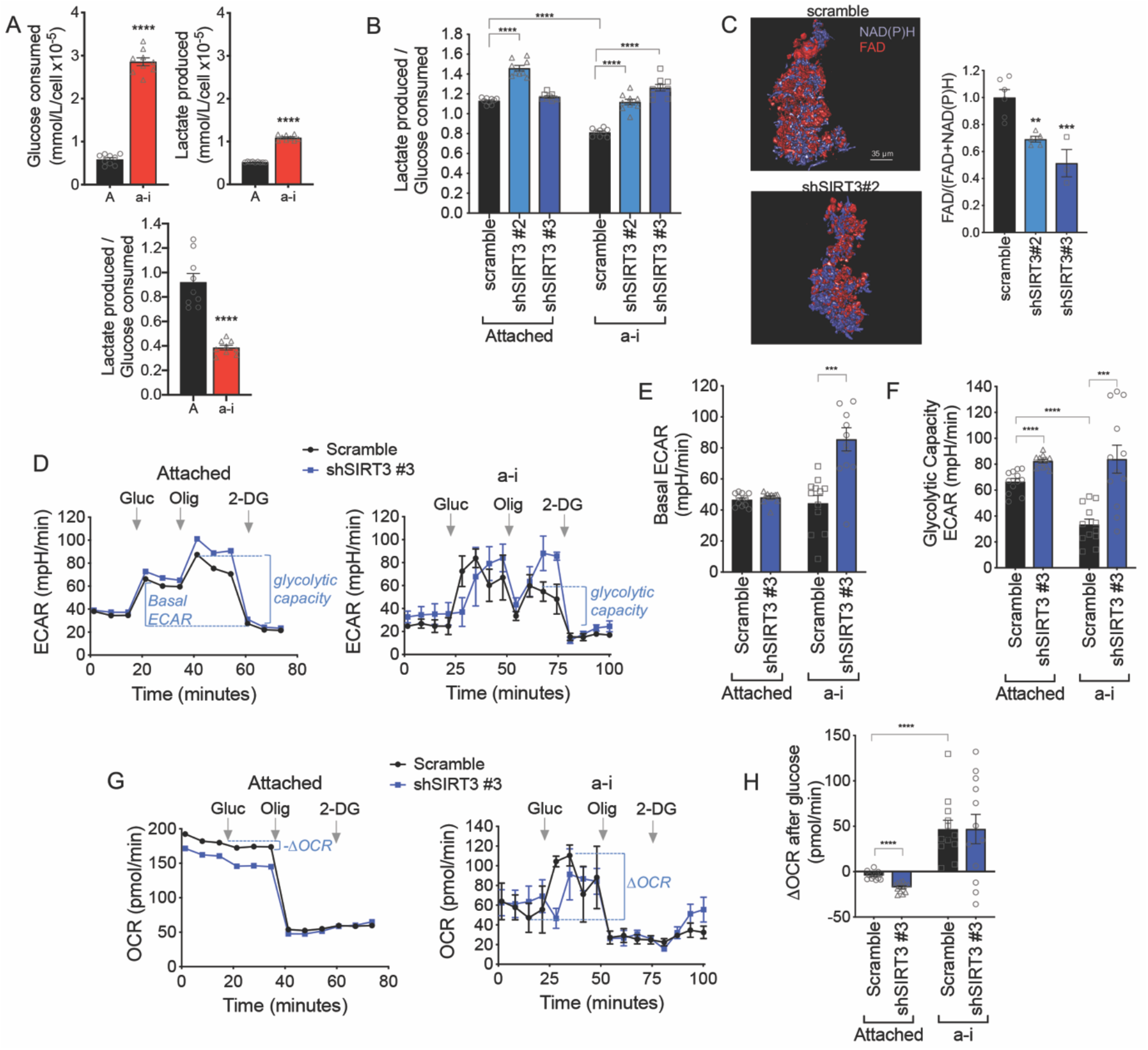
SIRT3 suppresses the glycolytic capacity of anchorage-independent (a-i) cells. **A.** OVCA433 a-i cultured cells (24 h) consume more glucose per cell compared to attached cells, but produce less lactate relative to glucose consumed (n=9 ± SEM; unpaired t-test ****P<0.0001). **B.** SIRT3 knock-down shifts OVCA433 cells towards enhanced lactate production relative to glucose consumption (n=9 ± SEM; one-way ANOVA P=0.01, Tukey’s multiple comparisons test ****P<0.0001). **C.** The optical redox ratio FAD / FAD + NAD(P)H is decreased in a-i conditions following SIRT3 knock-down in OVCA433 (n=3-5; unpaired t-test *P=0.04). **D.** Extracellular Acidification Rates (ECAR) were measured in attached and a-i OVCA433 cells using a Seahorse XFp extracellular flux analyzer. One representative experiment shown (n=3). 10mM glucose (Gluc), 1 μM Oligomycin A (Oligo) and 50mM 2-Deoxyglucose (2-DG) were added at indicated times. **E. & F.** SIRT3 knock-down increases basal ECAR/Glycolysis (E) and Glycolytic Capacity (F). **G.** Oxygen Consumption rate (OCR) was monitored simultaneously as in D. were determined as a change in ECAR or OCR following addition of Glucose, respectively. **H.** Attached cells display a decrease in OCR, while cells in a-i significantly increase their OCR following glucose addition. (E, F & H, n=12; ***P<0.001, *****P<0.0001)

As expected OCR was inhibited in response to glucose addition in attached conditions, as cells use glucose primarily for lactic acid production (**Fig. 3G**). Interestingly, extracellular flux analysis confirmed that detached cells do not utilize glucose primarily for lactate production, and also increase their oxygen consumption in response to glucose addition (**Fig. 3G&H**). While OCR was further suppressed by SIRT3 knock-down in attached cells, large variability in OCR readings between experimental replicates was unable to ascertain if SIRT3 directly inhibits Oxidative Phosphorylation in detached conditions.

### The SIRT3-dependent oxidant scavenging is necessary for ovarian cancer anchorage-independent survival

Since we found that SIRT3 is specifically increased upon cell detachment, we tested the necessity of SIRT3 for anchorage-independent survival. Knock-down of SIRT3 significantly increased the fraction of dead cells when OVCA433, SK-OV-3 and OVCAR3 cells were cultured in anchorage-independent conditions (**Fig. 4A, Supp. Fig. 3A**). This was accompanied by an inability of cells to rapidly aggregate into spheroid clusters within 6 hours following detachment (**Supp. Fig. 3B**), and resulted in loosely aggregated OVCA433 cells, and smaller and less uniform SK-OV-3 spheroids by 72 hours of anchorage-independence (**Fig. 4A**). Loss of SIRT3 expression increased the fraction of apoptotic cells following matrix detachment, while there was no effect of SIRT3 knock-down on apoptosis in attached conditions, suggesting that SIRT3 is inhibitory to anoikis (**Fig. 4B**). A pro-survival role for SIRT3 was also observed in single cell clonogenic assays. SIRT3 knock-down significantly inhibited colony number, but not average colony size, indicating that SIRT3 knock-down inhibits initial single cell survival and seeding, rather than proliferation (**Fig. 4C**). Accordingly, shRNAs to SIRT3 had no significant impact on cell cycle progression (**Fig. 4D, Supp. Fig. 3C**). Cells in detached conditions had fewer cells in S and G2/M phases of the cell cycle, suggesting that these cells slow their proliferation, as previously demonstrated [32], however this was unaffected by SIRT3 knock-down (**Fig. 4D**). These data demonstrate that SIRT3 is necessary for survival under anchorage-independence by inhibiting anoikis.

**Figure 4:**
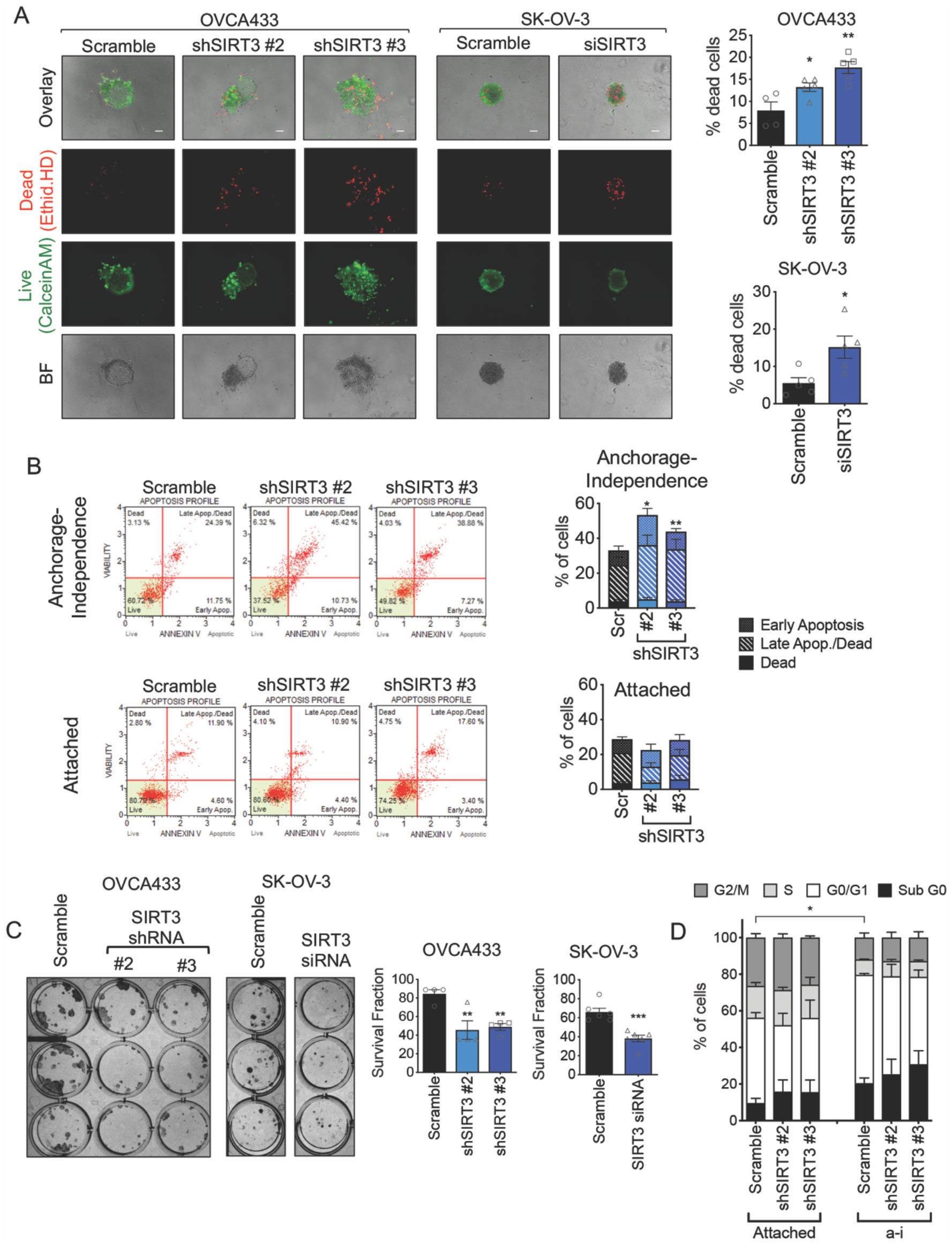
SIRT3 expression is required for anchorage-independent ovarian cancer cell survival. **A.** SIRT3 knock down increases the dead cell fraction of cells in anchorage-independent (a-i) spheroid aggregates, when cultured in ULA plates for 72 h. Cells were stained with Ethidium homodimer (dead cells) and Calcein AM (live cells) and fractions of live and dead cells quantified (n=4; OVCA433: one-way ANOVA P=0.002, Tukey’s multiple comparison test *P=0.04, **P=0.001; SK-OV-3: unpaired t-test *P=0.019; scale bar = 100μm). **B.** SIRT3 knock-down increases the apoptotic fraction of cells cultured for 24 h in anchorage-independence, but not in attached conditions. Apoptosis was assessed by Annexin V (n=4-5 experimental replicates; repeated measures ANOVA of total dead cell fraction, a-i: P=0.008, Attached: not significant; Bonferroni’s multiple comparison test *P<0.05, **P<0.01). **C.** SIRT3 knock-down inhibits single cell clonogenic survival (n=4-6; OVCA433: one-way ANOVA P=0.005, Tukey’s multiple comparison test **P<0.01; SK-OV-3: unpaired t-test *P=0.0004). **D.** SIRT3 knock-down does not significantly affect cell cycle progression in either attached or a-i cultured OVCA433 cells. (a-i, 24 h, n=3 experimental replicates; Two-way ANOVA, Tukey’s post-test *P=0.036, comparison of G0/G1).

As expected, SOD2, the target of SIRT3, was similarly necessary for anchorage-independent survival. SOD2 knock-down significantly increased the dead cell fraction and mitochondrial O_2^·^_^-^ levels of detached ovarian cancer cells (**Fig. 5A&B, Supp. Fig. 4**). Cell death and MitoSox fluorescence induced by SIRT3 knock-down was successfully rescued with a mitochondrial SOD2 porphyrin mimetic MnTBAP and the glutathione precursor N-acetyl-L-cysteine (NAC; **Fig. 5C&D, Supp. Fig. 5**), indicating that SIRT3-mediated oxidant scavenging is an important pro-survival mechanism for ovarian cancer cells following detachment.

**Figure 5:**
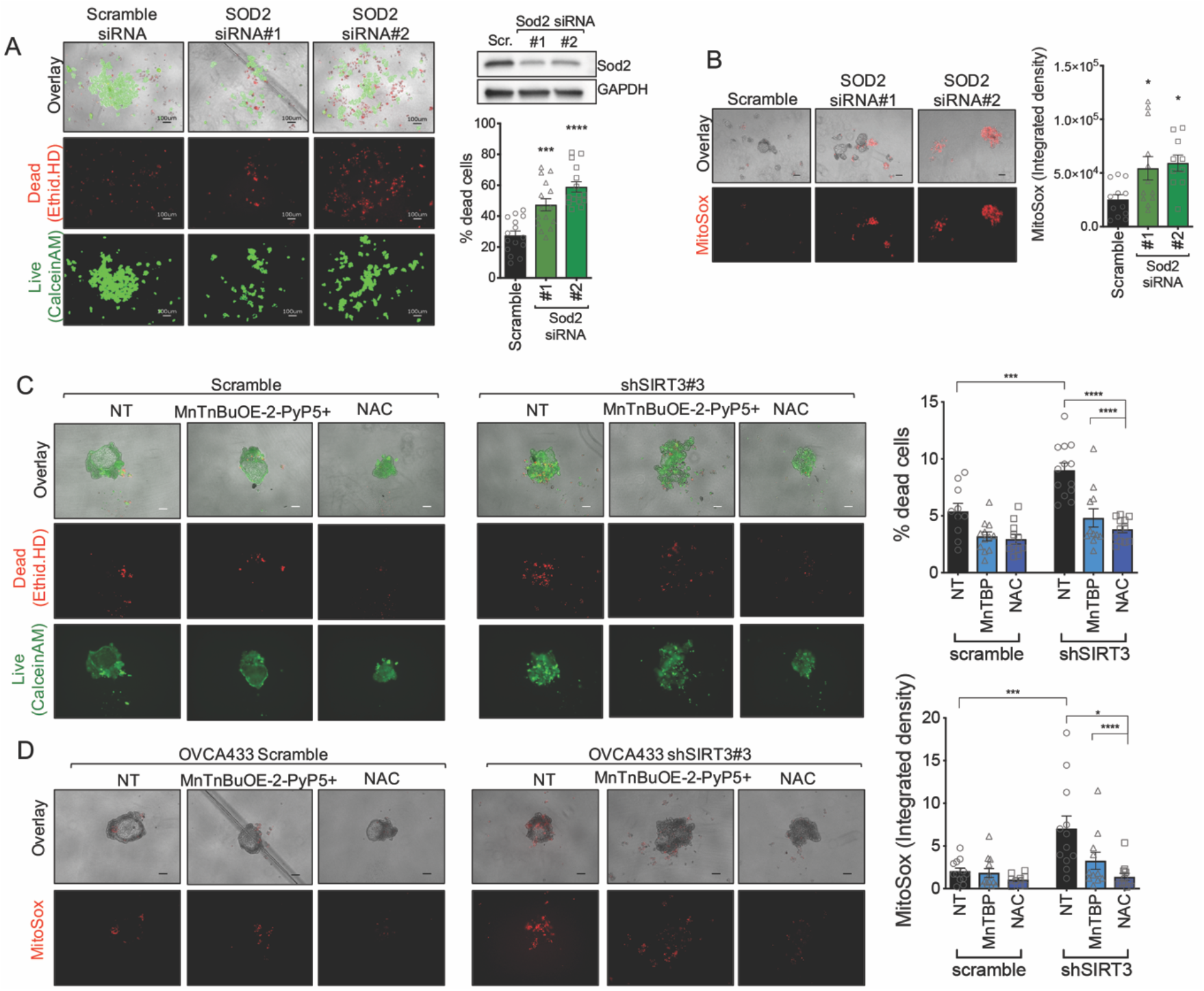
SOD2 and oxidant scavenging are required for anchorage-independent survival. **A.** SOD2 knock-down increases the dead cell fraction of OVCA433 cells in anchorage-independent (a-i) spheroid aggregates, when cultured in ULA plates for 72 h. Cells were stained with Ethidium homodimer (dead cells) and Calcein AM (live cells) and fractions of live and dead cells quantified (n=15; one-way ANOVA P<0.0001, Tukey’s multiple comparison test ***P=0.0003, ****P<0.0001). SOD2 expression was inhibited using siRNA in OVCA433 cells (Scr, scramble control) **B.** SOD2 knock-down increases oxidation and fluorescence of the mitochondrial O_2^·^_^-^ probe MitoSox in ULA cultured OVCA433 cells (n=10-12; one-way ANOVA P=0.01, Tukey’s multiple comparisons test *P<0.05). **C.** Co-treatment of cells with 10μM MnTBAP or 2mM NAC rescues OVCA433 cell viability following siRNA mediated SIRT3 knock-down; and **D.** results in decreased mitochondrial MitoSox oxidation (72 h a-i; n=12-15; ± SEM; OVCA433: one-way ANOVA P<0.0001, Tukey’s multiple comparisons test ****P<0.0001, **P=0.001; SK-OV-3: unpaired t-test ****P<0.0001).

### SIRT3 and SOD2 are necessary for metastatic colonization of the peritoneal cavity

Since anchorage-independent survival is a critical step for successful transcoelomic metastasis of ovarian cancer cells in the peritoneal cavity [3, 4], we tested if the increases in SIRT3 and SOD2 specific to detachment are necessary for peritoneal tumor formation *in vivo*. To transiently decrease SIRT3 and SOD2 expression during the anchorage-independent phase, SK-OV-3-luc cells were transfected with siRNAs targeting either SIRT3, SOD2 or a scramble control. Twenty-four hours later cells were detached and 1×10^6^ viable cells suspended in 150 μl PBS immediately injected into the peritoneal cavity of NSG mice (n=8), and tumor establishment monitored by bioluminescence imaging. Either SIRT3 or SOD2 knock-down significantly inhibited peritoneal tumor formation over time (**Fig. 6A-C**). Assessment of the omentum, a major target tissues of metastatic ovarian cancer [33], and a site where the majority of SK-OV-3 tumors were detected, revealed that control SK-OV-3 tumor cells replaced most of the adipocytes in this tissue. SIRT3 knock-down, and to a greater extent SOD2 knock-down, resulted in the establishment of fewer tumor nodules in the omentum, suggesting that the loss of these proteins leads to reduced seeding of viable tumor cells, likely as a consequence of increased anoikis during dissemination (**Fig. 6D-H**). The size of individual tumor nodules was not significantly different and varied widely in all experimental groups, although a trend in larger tumors was observed from control scramble siRNA transfected SK-OV-3 tumor cells (**Fig. 6H**). The above data demonstrate that SIRT3 and SOD2 are necessary for successful transcoelomic tumor formation *in vivo*.

**Figure 6:**
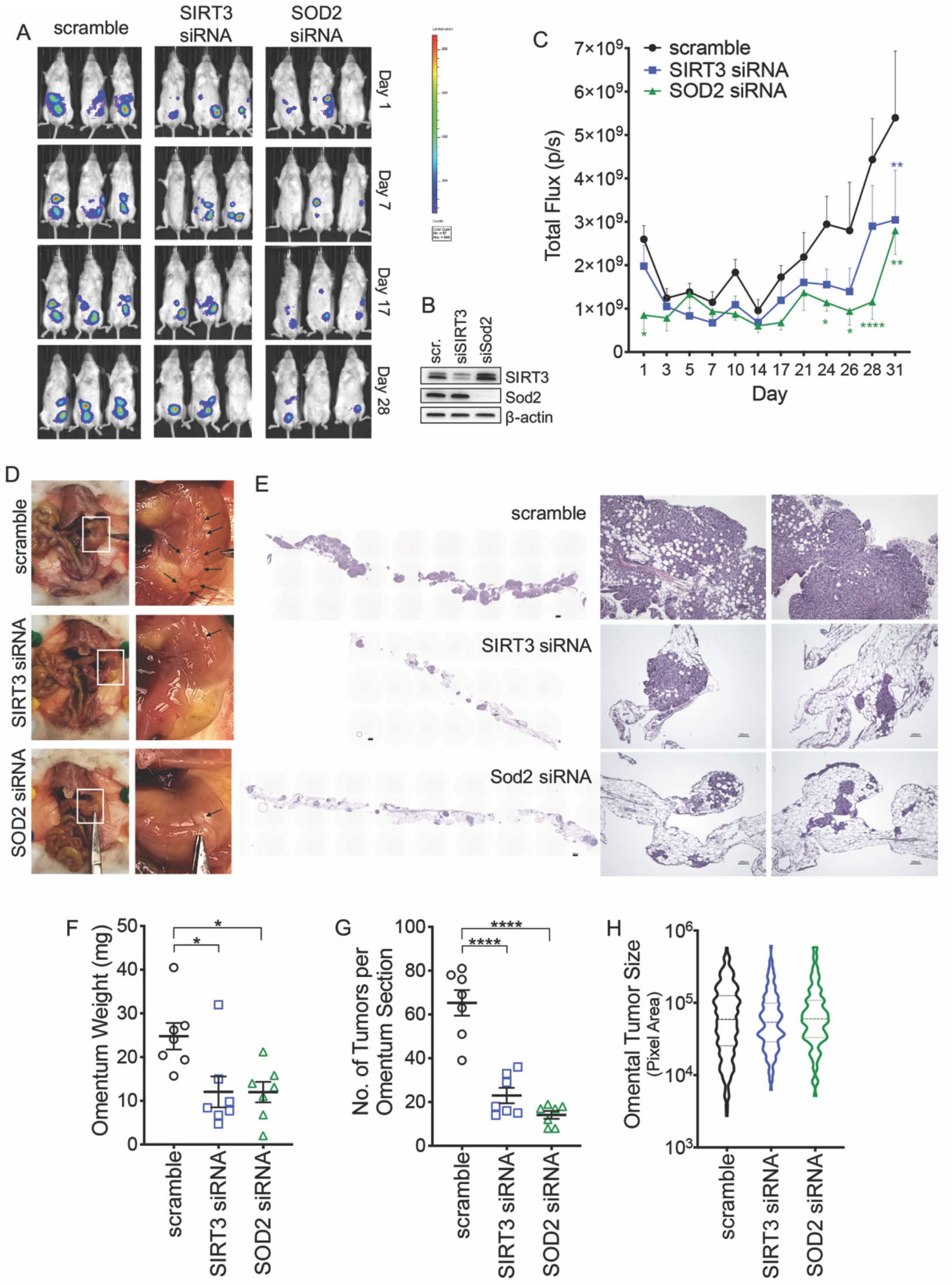
SIRT3 and SOD2 are required for successful metastasis to the omentum. **A.** Representative tumor luminescence images of NSG mice injected with SK-OV-3-luciferase cells transfected with either scramble siRNA, siRNA targeting SIRT3 or SOD2. **B.** Western blot demonstrating knock down of SIRT3 and SOD2 three days after transfection. **C.** Quantification of whole animal tumor luminescence over time (n=7-8; Mixed-design ANOVA P=0.004, Dunnett’s post test *P < 0.05, **P < 0.01, ****P < 0.0001). **D.** Assessment of the peritoneal cavity revealed that the majority of SK-OV-3 tumors (black arrows) were localized to the omentum. **E.** H & E staining of omental demonstrates that SIRT3 and SOD2 knock down abrogate tumor burden in the omentum. **F.** SIRT3 and SOD2 knock down decreases omental weight and **G.** number of tumors per longitudinal section of omentum assessed by H & E (One way ANOVA, F: P = 0.01, G: P < 0.0001; Tukey’s post test *P < 0.05, P < 0.0001). **H.** Spread of individual omental tumor sizes (violin plot, median + interquartile range).

## Discussion

Here, we present new evidence for the pro-metastatic role of SIRT3 in cancer, and demonstrate that the mitochondrial SIRT3/SOD2 stress response pathway is specifically upregulated in response to cellular detachment (**Fig. 1**), and specifically necessary for anchorage-independent survival and transcoelomic metastasis (**Fig. 4&6**).

Similar to several cancers previously reported [13–17], the expression of SIRT3 is low in primary tumors of high-grade serous adenocarcinoma samples from the Cancer Genome Atlas (TCGA). However, contrary to other tumor types, low SIRT3 expression was not associated with a decreased in overall patient survival (**Supp. Fig. 6**). The observation that a decrease in SIRT3 expression does not predict an unfavorable patient outcome somewhat contradicts previous reports of the anti-tumorigenic effects of SIRT3 in ovarian cancer cells [26, 27], and highlights that SIRT3 likely plays a dichotomous role during the progression of ovarian cancer. The context-dependent regulation of SIRT3 during ovarian cancer metastatic progression is evident by our findings that SIRT3 expression is significantly increased in cells derived from patient ascites, compared to matching primary ovarian tumor tissue (**Fig. 1**). We propose that this transient regulation addresses the seeming contradiction between SIRT3’s anti-proliferative properties and low primary tumor expression, and the role of SIRT3 as an important mitochondrial stress response gene important for cell survival.

Few studies have examined the context-specific regulation and function of SIRT3 in cancer. Most work demonstrating the anti-tumor role of SIRT3 was based on discoveries that SIRT3 expression is downregulated in a number of primary tumors, and that this decrease is associated with poor patient survival [13, 14, 16, 17]. The functional consequences of SIRT3 loss were mainly assessed in attached cell culture conditions or *in vivo* models testing the role of SIRT3 on primary tumor growth, without assessment of its function in metastatic progression [12, 18, 19]. However, our work and that of others is starting to unravel the complex role and regulation of SIRT3 during cancer progression. While we observed that manipulation of SIRT3 expression has few deleterious effects in attached cells, SIRT3 and its downstream target, the mitochondrial O_2^·^_^-^ scavenger SOD2 were required for survival following matrix detachment (**Fig. 4&5**), and this was necessary for successful peritoneal tumor formation *in vivo* (**Fig. 6**). Importantly, we found that the SIRT3-dependent increase in SOD2 activity is an early response to matrix detachment and sustained under long term anchorage-independence through SIRT3-mediated *SOD2* transcriptional regulation (**Fig. 2**), likely as a consequence of SIRT3-dependent FOXO3a deacetylation [25, 29]. Concurrent with our findings, SIRT3 has been implicated in anoikis resistance of oral squamous cell carcinoma cells [34], and for the maintenance of mitochondrial ROS scavenging in GBM stem cells [24]. In addition, the SIRT3/FOXO3a/SOD2 axis is upregulated as part of the mtUPR and necessary for breast cancer metastasis [25, 35]. Moreover, we and others have demonstrated that SOD2 is required by metastatic cells as an adaptation to oxidative stress, for the maintenance of mitochondrial fidelity, and as a regulator of mitochondria redox signaling [20, 36–38].

Highlighting the dichotomous role of SIRT3 in cancer is the finding that the SIRT3-dependent regulation of SOD2 is also a previously described mechanism of SIRT3’s anti-tumor activity. We find that mitochondrial oxidant scavenging is necessary for anchorage-independent survival, and that both SIRT3 and SOD2 are required for this (**Fig. 4&5**). Conversely, increased oxidant production in *Sirt3* knock-out mice was shown to induce tumor formation when *Sirt3−/−* MEFs were transformed with oncogenes Ras and Myc, as a result of increased DNA oxidation [12]. In addition, the anti-tumor effects of SIRT3 were demonstrated to be related to manipulation of tumor metabolism *via* SIRT3 inhibition of HIF-1α [18, 30]. The increase in mitochondrial O_2^·^_^-^ levels in SIRT3 knock-down cells was shown to result in HIF-1α protein stabilization as a consequence of prolyl-hydroxylase inhibition, and the increased glycolytic phenotype of SIRT3 knock-down cells associated with this ROS-mediated increase in HIF-1α-signaling [18, 30]. While we similarly found that SIRT3 suppresses glycolysis, increased HIF-1α stabilization with SIRT3 loss was a phenotype limited to attached conditions (**Supp. Fig. 7**). HIF-1α levels remained unaltered following SIRT3 knock-down in anchorage-independence, indicating that the effects of SIRT3 on glycolysis are not coupled to HIF-1α suppression in this state of tumor metastasis. Several other metabolic enzymes are known SIRT3 targets. For example, activation of IDH2 by SIRT3-dependent deacetylation can shift metabolism from glycolysis to oxidative phosphorylation [39], and the presence of IDH1 and IDH2 is important for survival of anchorage-independent cells by aiding in regeneration of NADPH for glutathione reduction [28]. Moreover, SIRT3 is a known activator of pyruvate dehydrogenase (PDH), resulting in enhance shuttling of pyruvate into the TCA cycle [40, 41]. It remains to be determined if these enzymes contribute to the suppression of glycolysis elicited by SIRT3 expression in anchorage-independence, or if alternate deacetylation targets of SIRT3 have functional roles in this stage of metastasis. As suggested by analysis of lactate levels and extracellular acidification rate measurements (**Fig. 3**), increased glucose uptake in anchorage-independent cells is more uncoupled from lactate production compared to attached cells. It is now well accepted that metabolic flexibility is a hallmark of cancer, allowing cells to cope with fluctuations in nutrient availability during different tumor stages and in response to changing tumor microenvironments. This is evident in anchorage-independent cancer spheroids, which are often enriched in cancer stem cells, and marked by the ability to readily switch their metabolism to induce the pentose phosphate pathway, TCA cycle and oxidative phosphorylation, based on nutrient availability and anabolic demands [6, 42]. As previously reported [32], we also observed a change in cell cycle in detached cells, suggesting that the switch from high lactate production to glucose utilization for other metabolic pathways may be a reflection of a switch from pro-proliferative energy demands in attached cells to pro-survival metabolism in anchorage-independence. The observed suppression of glycolytic capacity by SIRT3 in detached cells requires further attention, and could be an important mechanism for cancer cells to survive metastatic spread in new tumor environments with alternate carbon sources, including the ascites fluid and the adipocyte-rich omentum [43]. In addition, the suppression of glycolytic flux by SIRT3 may be a further survival mechanism to prevent lactic acid toxicity.

In conclusion, our data suggest that the rapid and sustained upregulation of SIRT3 following matrix detachment is necessary for SOD2-mediated mitochondrial oxidant scavenging to enhance anchorage-independent survival and peritoneal colonization of ovarian cancer cells. Additional consequences of the context specific increase of SIRT3 on metabolic changes likely further aid in transcoelomic spread as a response to changing nutrient environments. Our study highlights the context-dependent role and regulation of SIRT3 in cancer and has important implications for future targeting of this protein for anti-cancer therapies.

## Materials and Methods

### GEO data set

GEO data set GSE85296 was examined using GEO2R (NCBI) to determine SIRT3 and SOD2 expression in matching ovarian, peritoneal, omental and ascites specimens from four ovarian serous adenocarcinoma patients.

### Cell lines and cell culture conditions

OVCA433 and OVCA420 cells were provided by Dr. S.K. Murphy (Duke University). ES-2 and NIH-OVCAR3 cells were purchased from American Type Culture Collection (CRL-1978, HTB-161; ATCC Manassas, VA). OVCA433, OVCA420 and OVCAR3 cells were cultured in RPMI media supplemented with 10% FBS. Luciferase expressing SK-OV-3-Luc cells were from the Japanese Collection of Research Bioresources Cell Bank and cultured in McCoy’s 5A medium with 10% FBS. All cells were maintained at 37°C with 5% CO_2_. Cell authentication was carried out by the commercial provider or in-house through STR genotyping. In rescue experiments cells were co-treated with the Manganese Porphyrin O_2_^•−^ scavenger ortho tetrakis(N-n-butoxyethylpyridinium-2-yl) porphyrin (MnTnBuOE-2-PyP5+), provided by Dr. Ines Batinic Haberle (Duke University), or with N-acetyl-L-cysteine (Sigma), for indicated times and doses.

### Culturing cells in ULA plates

Cells were trypsinized, counted, and equal viable cells seeded in 6-well Ultra Low Attachment (ULA; Corning) plates (1×10^5^ cells/2 mL/well) for RNA and protein extraction. For live/dead and MitoSOX staining, cells were seeded 1×10^3^ cells/200 μL/well in 96-well round-bottom ULA plates.

### EOC specimens from patient ascites

Ascites specimens from patients diagnosed with high grade serous adenocarcinoma (Stage III-IV) were collected at the Penn State Cancer Institute (Hershey, PA) and the London Health Sciences Centre (London, Ontario), with approval granted from the Penn State College of Medicine and Western University Ontario institutional research ethics boards, respectively. EOCs were isolated from ascites, as previously described [44], and maintained in culture at 37°C, 5% CO_2_ in MCDB/M199 medium supplemented with 10% FBS and penicillin/streptomycin. Microarray analysis was carried out on total RNA extracted from EOCs obtained at Western University, as previously described [45], using the Ad-GFP transduced controls. RNA was extracted following culturing for 72 h in ULA plates or from controls grown in attached conditions. Affymetrix Human Genome U133A GeneChip analysis (Santa Clara, CA) was carried out at Precision Biomarker Resources Inc. (Evanston, IL).

### Semi-quantitative real-time RT-PCR

RNA was extracted using the Direct-zol RNA MicroPrep (Zymo Research, R2062) and cDNA synthesized using the iScript cDNA kit (Bio-Rad, 1708891). Real-time RT-PCR was performed using the iTaq Universal SYBR Green Supermix (Bio-Rad) and Bio-Rad CFX96 Real-Time PCR System. Relative SIRT3 mRNA expression was normalized to the geometric mean of 4 housekeeping genes: HPRT1, TBP, 18S, and GAPDH, and data analyzed using the ΔΔCt method. The following primers were used: SIRT3-sense (S): 5’-AGCCCTCTTCATGTTCCGAAGTGT-3’; SIRT3-antisense (AS): 5’-TCATGTCAACACCTGCAGTCCCTT-3’; HPRT1-S: 5’-TGACCTTGATTTATTTTGCATACC-3’; HPRT1-AS: 5’-CGAGCAAGACGTTCAGTCCT-3’; TBP-S: 5’-TTGGGTTTTCCAGCTAAGTTCT-3’; TBP-AS: 5’-CCAGGAAATAACTCTGGCTCA-3’; 18S-S: 5’-AGAAACGGCTACCACATCCA-3’; 18S-AS: 5’-CACCAGACTTGCCCTCCA-3’ GAPDH-S: 5’-GAGTCAACGGATTTGGTCGT-3’; GAPDH-AS: 5’-TTGATTTTGGAGGGATCTCG-3’. SOD2-S: 5’-TCCACTGCAAGGAACAACAG-3’ SOD2-AS: 5’-CGTGGTTTACTTTTTGCAAGC-3’.

### RNA interference

Short hairpin RNA (shRNA) with non-targeting scramble sequence (5’-GCACTACCAGAGCTAACTCAGATAGTACT-3’) or targeting SIRT3 sequences (shSIRT3_#1: 5’-GTACAGCAACCTCCAGCAGTACGATCTCC-3’; shSIRT3_#2: 5’-AACCAGAATATGTGAACTGAGTGGACACC-3’; shSIRT3_#3: 5’-TCTTCACTCTGCTGAAGCTCCTAATGGAA-3’; shSIRT3_#4: 5’-TCACATTCTGTTGACTCTCCATACTCAGC-3’) in pRS vector (Origene, TR309432) were used to stably transfect OVCA433 cells (**Supp. Fig. 1A**). Clones expressing shSIRT3_#2 and shSIRT3_#3 were used for all subsequent experiments. Scramble non-targeting SMARTpool control (D-001810-10-05), SIRT3-specific SMARTpool siRNA oligonuecliotides (L-004827-01-0005), SOD2-specific siRNA oligonucleotides (siSOD2_#1: 5’-AAGUAAACCACGAUCGUUA-3’, J-009784-06-0005 and siSOD2_#2: 5’-CAACAGGCCUUAUUCCACU-3’, J-009784-08-0005) were obtained from Life Technologies and 60pmol/1×10^5^ cells transfected using Lipofectamine RNAiMax.

### Immunoblotting

Lysates were prepared in RIPA buffer supplemented with protease and phosphatase inhibitors, and equal protein separated on tris-glycine SDS-PAGE, followed by transfer to PVDF membranes. Membranes were blocked in 5% milk/TBS/0.1%Tween-20, and incubated overnight at 4°C with the following antibodies: SIRT3 (5490S, Cell Signaling Technology); Histone H3 (9715S, Cell Signaling Technology); SOD2 (ab13533, Abcam); acetyl(K68)-SOD2 (ab137037, Abcam); GAPDH (AM4300, Invitrogen); β-actin (AM4302, Invitrogen); HIF1α antibody (610958, BD). HRP-conjugated secondary antibodies were obtained from GE Healthcare. Blots were visualized on a ChemiDoc MP system (Bio-Rad) using Femto ECL chemiluminescence substrate (Thermo Scientific).

### Live/dead staining

Cell viability was determined by staining cells with 2μM Calcein AM and 4μM ethidium homodimer (Sigma) in PBS for 30 min at 37 °C, followed by imaging on a Keyence BZ-X700 fluorescence microscope and analysis using Image J.

### Annexin V and Cell cycle Analysis

Apoptosis Annexin V and Cell Cycle analysis were performed using a Muse Flow cytometer (Sigma Milipore), according to manufacturer’s instructions.

### Clonogenicity assay

Single cell survival clonogenicity assays were performed as previously described [46, 47]. Briefly, 45 cells/well were plated in 12-well cell culture dishes and colonies (≥50 cells/colony) visualizing with 0.05% crystal violet dye.

### Mitochondrial Superoxide O_2^·^_^-^ Detection

5 μM MitoSOX Red (Invitrogen) was added to cells in HBSS for 30 min followed by washing. Images were captured using a Keyence BZ-X700 fluorescence microscope and fluorescence signal quantified using Image J.

### SOD Zymography

SOD activity was analyzed using zymography, as previously described [48, 49]. Briefly, cells were lyzed by sonication in potassium phosphate buffer (pH 7.8, 0.1 mM EDTA), and 50 μg of the proteins resolved by electrophoresis in a 10% non-denaturing polyacrylamide gel. Gels were incubated for 15 min in the dark with 2.5mM nitro blue tetrazolium, 30 mM TEMED, 0.028 mM riboflavine, 50 mM phosphate buffer, pH7.8, followed by washing in H_2_O, and visualization of SOD activity by light exposure.

### Media Glucose and Lactate Quantification

Cells were seeded in 6-well adherent or ULA plates (1×10^5^ cells/2 mL/well). After 24 h, media were collected by centrifugation and 200 μL analyzed for glucose and lactate content using the YSI 2900D Biochemistry analyzer (Xylem, Yellow Springs, OH), and values corrected for cell numbers.

### Optical Redox Ratio imaging

Anchorage-independent spheroids were imaged using the Nikon A1 MP+ Multi-Photon Microscope system (Nikon Instruments, New York) to assess endogenous fluorescence signals produced by NAD(P)H and FAD, as detailed in Supplemental Methods.

### Extracellular flux analysis

Oxygen consumption rate (OCR) and extracellular acidification rate (ECAR) were measured using a Seahorse XFp Analyzer (Agilent). 24 h prior to the assay cells were seeded at a density of 10,000 cells well into a XFp cell culture plate (attached conditions). Anchorage-independent cells were cultured for 24 h in 96 well round bottom ULA plates, followed by washing in assay media, and transfer of 10 spheroids into each assay well (1,000 cells per spheroid). Basal glycolysis was derived as the difference between ECAR following addition of glucose (10mM) and inhibition of glycolysis by 2-deoxy-glucose (50mM). Glycolytic capacity/maximal ECAR was determined following addition of the mitochondrial ATP synthase inhibitor Oligomycin (1μM).

### Intraperitoneal *in vivo* xenografts

SK-OV-3-Luc cells were transfected with scramble, SOD2 or SIRT3 siRNA (Dharmacon, as above). Cells were detached 24 h after transfection, washed, counted and 1×10^6^ viable cells resuspended in 150 μl of PBS before immediate IP injection into female Nod *scid* gamma mice (NSG, bred in house). Approval for animal studies was sought from the Penn State College of Medicine IACUC prior to study commencement. Luminescence imaging carried out every 2-3 days using an IVIS luminescence imaging system 10 min after mice were injected with 10 μl/g of body weight 15 mg/mL *in vivo* grade D-Luciferin (PerkinElmer). Mice were sacrificed by CO_2_ asphyxiation followed by cervical dislocation if they reached AAALAC-defined endpoints, or at day 31 post tumor cell injection. At necropsy organs were preserved in 10% buffered formalin, followed by paraffin embedding, sectioning, nd staining with hematoxylin and eosin. Number and size of tumors per longitudinal section of each omentum were imaged by microscopy and quantified in a blinded manner using Image J.

### Statistical analysis

All data are representatives of at least three independent experiments, unless otherwise noted. Data are expressed as mean ± SEM, and statistical analysis performed using GraphPad Prism Software v8, with statistical tests chosen based on experimental design, as described in figure legends.

## Acknowledgements

We thank Usawadee Dier, Dr. LP Madhubhani Hemachandra, Dr. Sarah Engelberth, Larissa Suparmanto, Sadie Dierschke, and Gina Deiter for technical assistance. We thank Dr. Arati Sharma and Dr. David Claxton (Penn State) for providing PDX animals. We are grateful for Dr. Vonn Walter’s advice on TCGA data analysis. MnTnBuOE-2-PyP5+ was kindly provided by Dr. Ines Batinic Haberle (Duke University). OVCA433 and OVCA420 cells were a generous gift from Dr. Susan Murphy (Duke University). This work was supported by NIH grants R00CA143229 (N.H.), R01CA230628 (N.H. & M.K.), S100D018124 (T.A.), by the Rivkin Center for Ovarian Cancer (NH), an equipment grant from Seahorse/Agilent (N.H.), and the Penn State Cancer Institute Developmental Fund Award (N.H.). Collection of ascites was partially supported by DoD Pilot award W81XWH-16-1-0117 (N.H.).

## Conflict of Interest

The authors declare no competing financial interests in relation to the work described.

## Author Contributions

Y.S.K. and P.G.V. contributed to study design, manuscript writing, and the majority of experimental execution and data analysis. V.M.J. and D.H.S. assisted with cell culture studies. L.C.C. carried out Seahorse experiments. B.L.W. and S.S. contributed to *in vivo* studies and data analysis. C.W.C. and K.M.A. performed YSI experiments. T.A. assisted in multiphoton experiments and performed data analysis. T.G.S. carried out microarray expression and data analysis. J.I.W. performed analysis of tumor sections. N.Y.L. assisted in data interpretation and manuscript editing. R.P. provided patient ascites and tumor cells and assisted in study design. K.M. contributed to conceptual design, data interpretation and writing of the manuscript. N.H. conceived and supervised the study, designed experiments and wrote the manuscript.

## Supplemental Figures

**Supplemental Figure 1:**
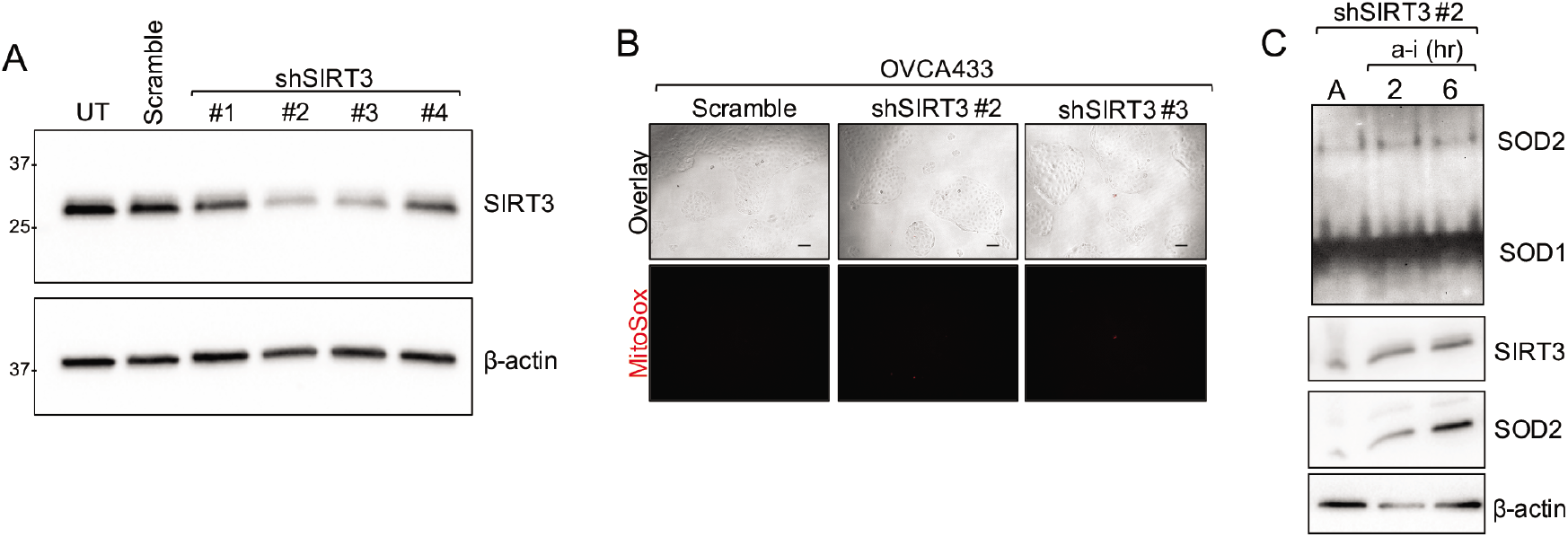
**A** SIRT3 protein expression knock-down following transient transfection of four shRNAs targeting SIRT3. shRNAs #2 and #3 were chosen for the establishment of OVCA433 stable knock-down cells. **B.** SIRT3 knock-down has little effect on superoxide levels in attached culture conditions, as assessed by MitoSox staining. **C.** SOD2 activity is inhibited by SIRT3 knock-down in early time points of matrix detachment. SOD2 activity was assessed by zymography in attached (A) OVCA433 cells and cells cultured for 2 or 6 h in anchorage-independence (a-i).

**Supplemental Figure 2:**
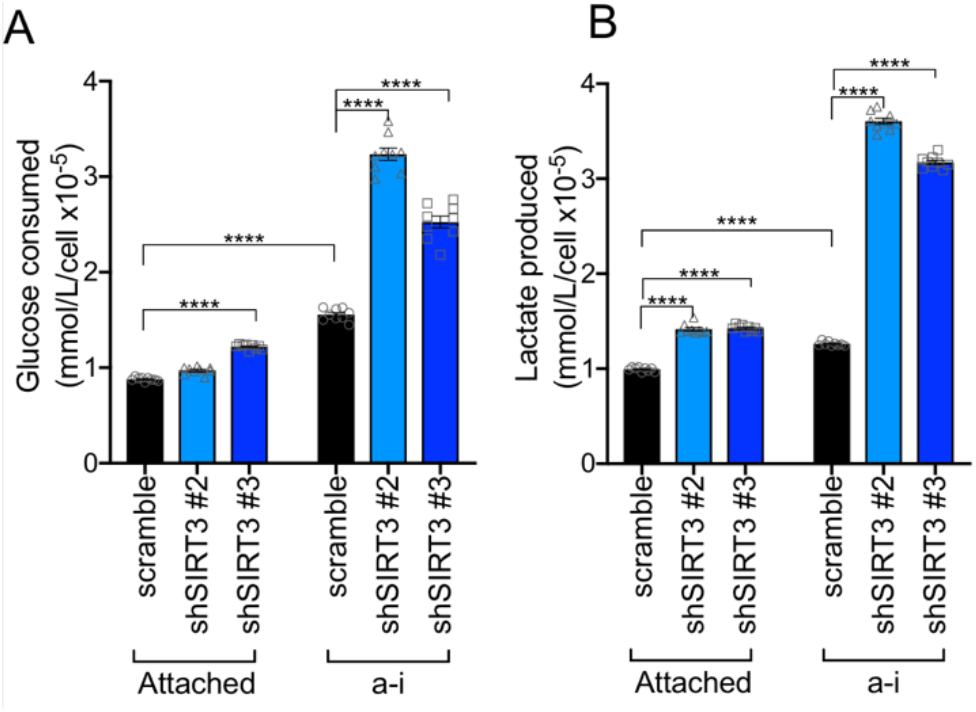
Media glucose (**A**) and lactate (**B**) levels were measured using the YSI biochemical analyzer and expressed as Glucose consumed and lactate produced by correcting for media glucose and lactate levels respectively, and expressed relative to cell numbers (n=9; one-way ANOVA P=0.01, Tukey’s multiple comparisons test ****P<0.0001).

**Supplemental Figure 3:**
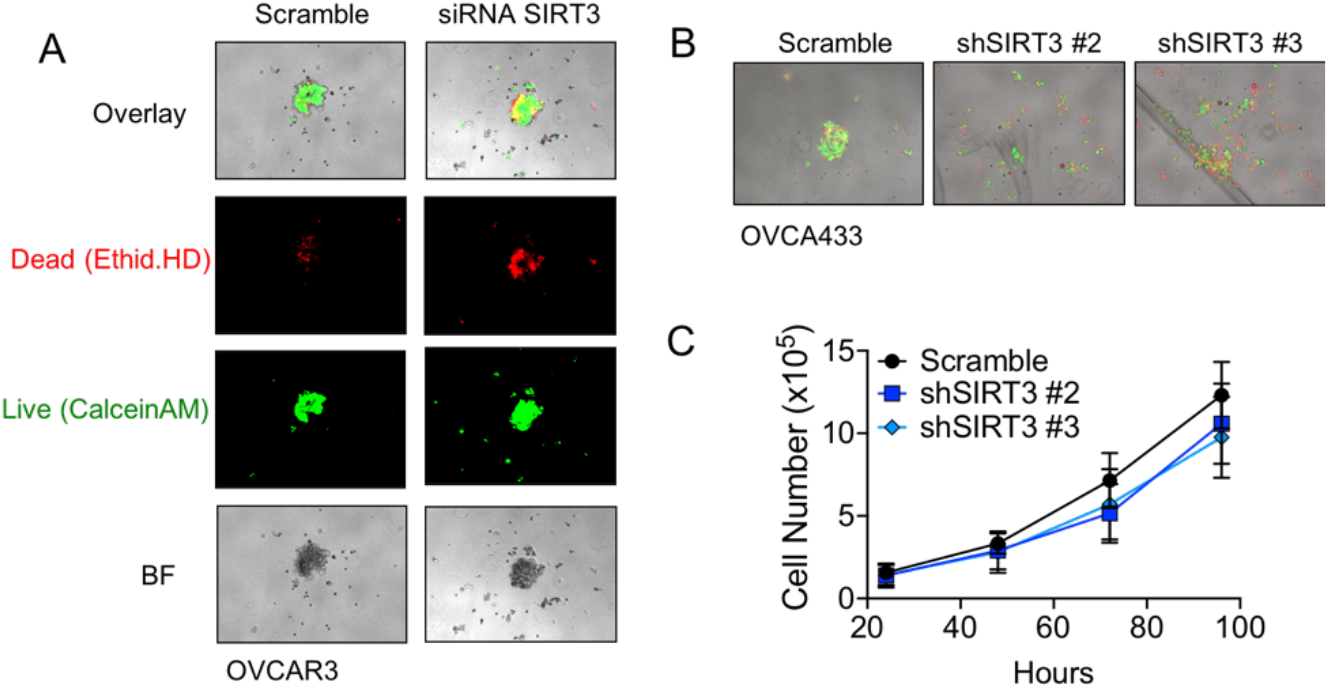
**A.** SIRT3 knock-down via siRNA delivery increases the dead cell fraction of OVCAR3 cells cultured as spheroid aggregates for 72 h in ULA plates. Cells were stained for live and dead cells using Ethidium HomoDimer and Calcein AM, respectively. **B.** Stable SIRT3 knock-down via shRNA inhibits rapid aggregation of OVCA433 in anchorage-independent cell culture conditions. 1,000 cells were plated per well in 96 well ULA culture plates and live dead staining carried out 6 hours after seeding. **C.** SIRT3 knock-down does not significantly affect OVCA433 cell proliferation in attached conditions (n=3).

**Supplemental Figure 4:**
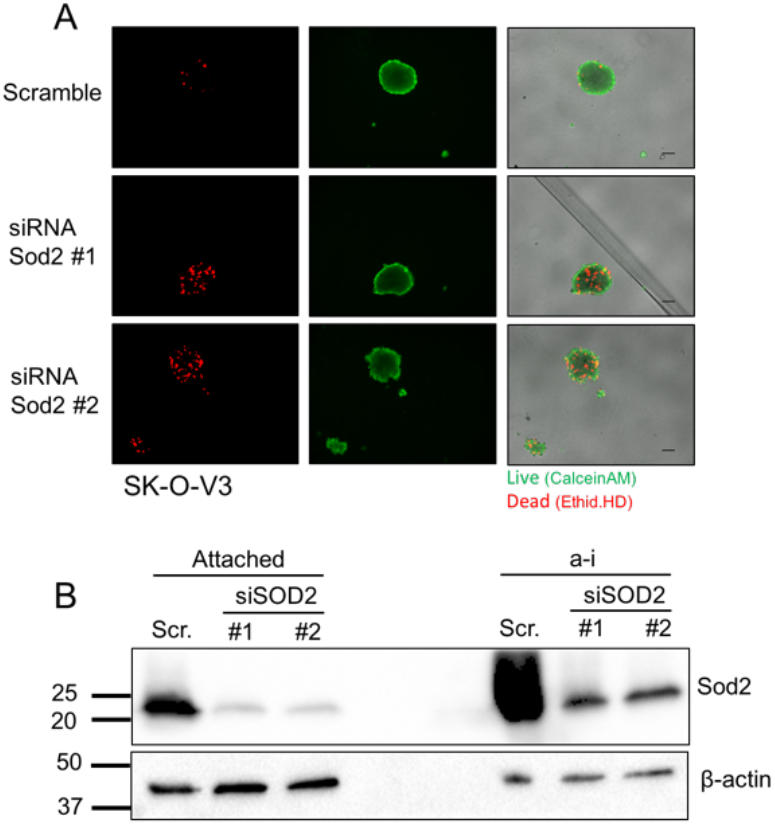
**A.** SOD2 knock-down decreases cell viability in SK-OV-3 cells cultured in anchorage-independence for 72 h. Cells were stained for live and dead cells using Ethidium HomoDimer and Calcein AM, respectively. **B.** Western blot analysis of Sod2 expression in SK-OV-3 cells following siRNA mediated knock-down.

**Supplemental Figure 5:**
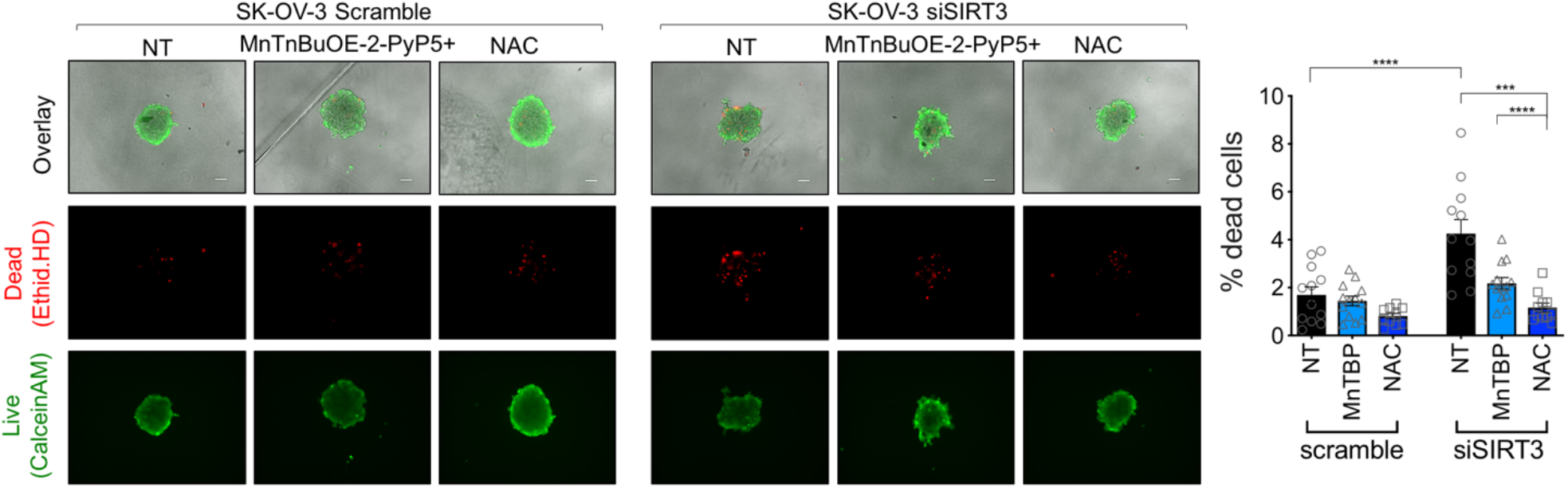
Treatment of cells with 10μM MnTBAP or 2mM NAC rescues SK-OV-3 cell viability.

**Supplemental Figure 6:**
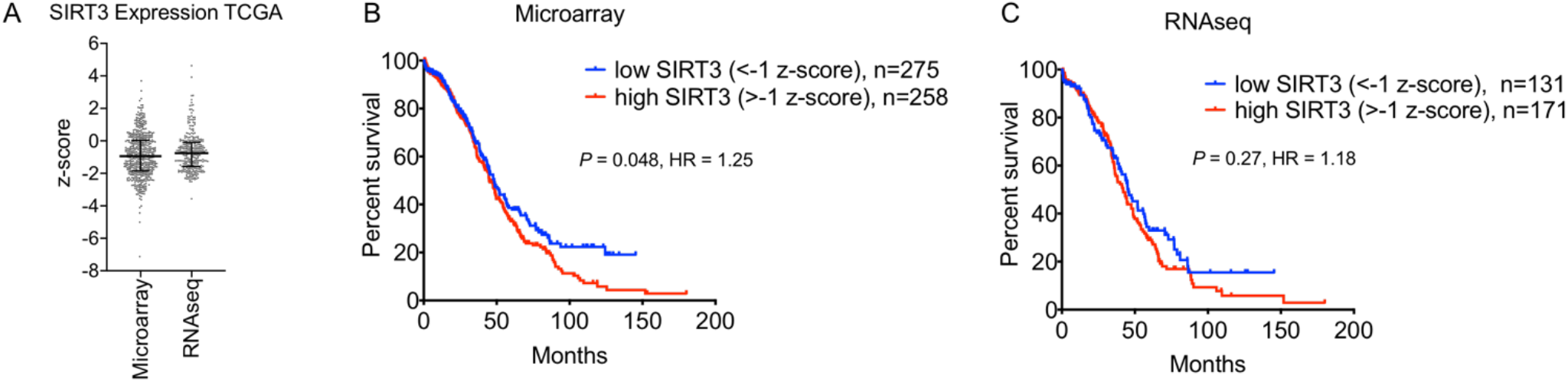
**A.** Spread of SIRT3 expression in TCGA serous ovarian adenocarcinoma samples subjected to Agilent micro-array (n=533) and RNAseq (n=303) analysis (median with interquartile range indicated). **B & C.** Kaplan Meier curves of overall survival of samples subjected to Microarray analysis (**B**), and RNA seq analysis (**C**). Samples with SIRT3 expression z-score <−1 were compared to those with z score >−1. (Log-rank Mantel-Cox test).

**Supplemental Figure 7:**
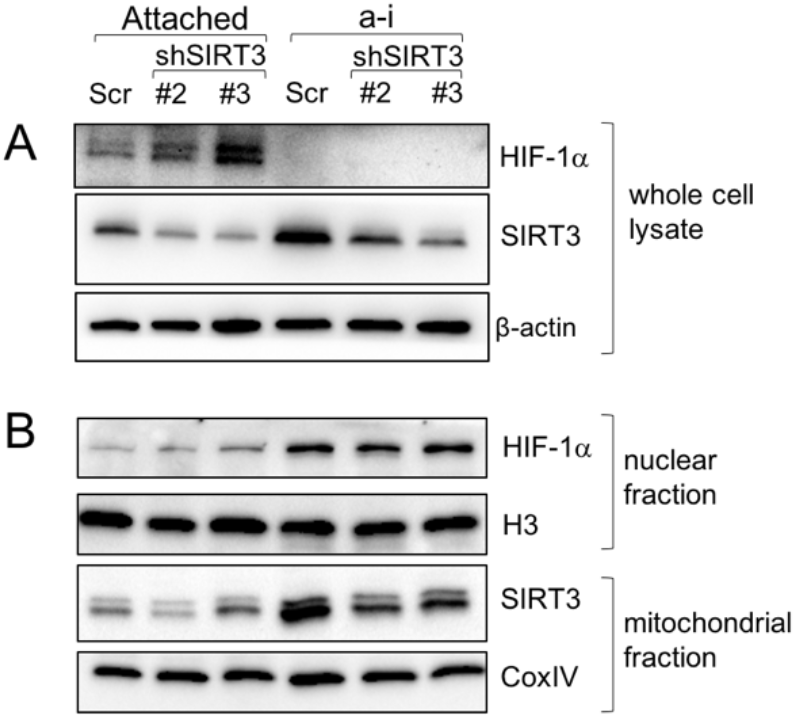
SIRT3 knock-down does not affect HIF-1α levels in anchorage-independent conditions (a-i). **A.** SIRT3 knock-down increases total HIF-1α levels in attached conditions only. (Representative blot shown - same lysates as Figure 2F). **B.** HIF-1α nuclear localization is enhanced in a-i compared to attached conditons. SIRT3 knock-down does not affect HIF-1α nuclear localization in a-i.

### Supplemental Methods

#### Optical Redox Ratio imaging

Anchorage-independent spheroids were imaged using the Nikon A1 MP+ Multi-Photon Microscope system (Nikon Instruments, New York) to assess endogenous fluorescence signals produced by NAD(P)H and FAD, using a mode-locked femto-second Spectra-Physics InSight DS femtosecond single-box laser system with automated dispersion compensation tunable between 680-1300 nm (Spectra-Physics, Mountain View, CA) and Nikon scan head coupled with Nikon upright microscope system (Nikon Instruments, New York). The laser output was attenuated using AOTF and the average power was consistently maintained below the damage threshold of the samples. The laser beam tuned to 1000 nm was then focused on the specimen through a high numerical aperture, low magnification, long working distance, dipping objective, CFI75 Apo Water 25X/1.1 LWD 2.0mm WD, and backscattered emissions collected through the same objective lens. Nikon Element Software was used for the image acquisition. In the reflection mode, non-descanned high-sensitivity GaAsP detectors were used for very efficient signal detection. A 750 nm Dichroic was used to prevent the scattered IR laser radiation from reaching the detector and a 460 nm long pass dichroic beam splitter (460 DCLP, Chroma Technology, USA) was used to collect NAD(P)H signal below 460 nm, a 560 nm long pass dichroic beam splitter (560 DCLP, Chroma Technology, USA) and a 660 nm long pass dichroic beam splitter (660 DCLP, Chroma Technology, USA) were used to collect FAD signal above 560 and below 660 nm. Spectral measurements to confirm the presence of NAD(P)H and FAD signals were also performed using 32-channel Nikon Spectral Detector integrated with Nikon A1 MP+ Multi-Photon Microscope system. For 3D image data set acquisition, the multiphoton excitation beam tuned to 1000 nm was first focused at the maximum signal intensity focal position within the spheroid sample and the appropriate detector levels were then selected to obtain the voxel intensities within range of 0-4095 (12-bit images) using a color gradient function. Later on, the beginning and end of the 3D stack (i.e. the top and the bottom optical sections) were set based on the signal level degradation. A series of 2D Images for a selected 3D stack volume were then acquired at 512 × 512 pixels. The 3D stack images with optical section thickness (z-axis) of approximately 2.0 μm were captured with voxel size of 0.5 × 0.5 × 2μm. For each spheroid volume reported, z-section images were compiled and the 3D image restoration performed using VOLOCITY (Perkin Elmar, UK). The volume estimation was performed on the 3D image data sets recorded from at least 3 spheroids. A noise removal filter was applied, and the lower threshold level in the histogram set to exclude all possible background voxel values. Sum of all voxels intensities above this threshold level was determined to be total NAD(P)H and FAD signals. The optical redox ratio was calculated from mean voxel intensity values using the equation, FAD/(FAD+NAD(P)H).

#### TCGA analysis

Agilent micro-array (n=533) and RNAseq (n=303) expression data from high grade serous adenocarcinomas were obtained from the cancer genome atlas (TCGA), using the cBioPortal interface (cBioportal.org; z-score spread of SIRT3 expression) ^1^. Overall survival was plotted using GraphPad Prism software and statistical differences determined using log rank test (Mantel-Cox).

## Notes

**Financial Support:** This work was supported by NIH grants R00CA143229 (N.H.), R01CA230628 (N.H. & M.K.), S100D018124 (T.A.), by the Rivkin Center for Ovarian Cancer (NH), an equipment grant from Agilent (N.H.), and the Penn State Cancer Institute Developmental Fund Award (N.H.). Collection of ascites was partially supported by DoD Pilot award W81XWH-16-1-0117 (N.H.).

